# Cell cycle dynamics of human pluripotent stem cells primed for differentiation

**DOI:** 10.1101/546291

**Authors:** Anna Shcherbina, Jingling Li, Cyndhavi Narayanan, William Greenleaf, Anshul Kundaje, Sundari Chetty

## Abstract

Understanding the molecular properties of the cell cycle of human pluripotent stem cells (hPSCs) is critical for effectively promoting differentiation. Here, we use the Fluorescence Ubiquitin Cell Cycle Indicator (FUCCI) system adapted into hPSCs and perform RNA-sequencing on cell cycle sorted hPSCs primed and unprimed for differentiation. Gene expression patterns of signaling factors and developmental regulators change in a cell cycle-specific manner in cells primed for differentiation without altering genes associated with pluripotency. Furthermore, we identify an important role for PI3K signaling in regulating the early transitory states of hPSCs towards differentiation.

## INTRODUCTION

Despite recent advances in generating specialized cell types from human pluripotent stem cells (hPSCs), many studies have noted that pluripotent stem cell lines often have an inherent inability to differentiate even when stimulated with a proper set of signals^1–5^. The cell cycle, particularly the G1 phase, may play an important role in enhancing the differentiation potential of PSCs^2,6–9^. However, simply lengthening the G1 phase in embryonic stem cells is not sufficient to facilitate differentiation^10^, suggesting that an improved understanding of the molecular properties of the embryonic cell cycle is needed.

In a prior study, we demonstrated that transiently treating hPSCs with dimethylsulfoxide (DMSO) for 24h prior to directed differentiation significantly increases the propensity for differentiation across all germ layers. This technique is now used by multiple laboratories to improve differentiation across species (including mouse, rabbit, primate, and human) into more than a dozen lineages, ranging from neurons and cortical spheroids to smooth muscle cells to hepatocytes^11–23^. While the DMSO treatment activates the retinoblastoma protein (Rb) and increases the percentage of hPSCs in the G1 phase of the cell cycle^2,24^, it remains unknown whether the DMSO treatment simply enriches cells in G1 or whether there are intrinsic changes to the cell cycle following the DMSO treatment that may potentiate differentiation.

Here, we use Fluorescence Ubiquitin Cell Cycle Indicator (FUCCI) technology to systematically track and understand cell division in hPSCs primed and unprimed for differentiation. The FUCCI system fuses red- and green-emitting fluorescent proteins to the cell cycle ubiquitination oscillators, Cdt1 and Geminin, whereby cdt1 tagged with RFP is present only when cells are in G1 and geminin tagged with GFP is only present when cells reside in the S/G2/M phases^25^. By performing RNA-sequencing on hPSCs sorted from the early G1, late G1, and SG2M phases of the cell cycle, we show that gene expression patterns of signaling factors and developmental regulators change in a cell cycle-specific manner in cells primed for differentiation following a 24h DMSO treatment. Changes in signaling pathways controlling cell proliferation, differentiation, and apoptosis, particularly the phosphoinositide 3-kinase (PI3K) pathway, were regulated by the DMSO treatment. Concordantly, transiently inhibiting PI3K signaling enhances hPSC differentiation across all germ layers.

To our knowledge, this is the first study to systematically perform RNA-seq on cell cycle sorted populations of hPSCs to investigate changes that occur within a cell line as cells transition towards a state for differentiation. This comprehensive analysis begins to shed light on important signaling pathways, particularly PI3K, in regulating the developmental potential of hPSCs during early transitory states.

## RESULTS

### Gene expression dynamics associated with cell cycle progression in hPSCs

To begin, we use the FUCCI system adapted into the hPSC H9 cell line^7^ to systematically track and isolate hPSCs from different phases of the cell cycle (Figure 1A). H9 FUCCI hPSCs were cultured under maintenance conditions in mTESR (control) or with 2% DMSO for 24h to prime hPSCs for differentiation (Figure 1B). Following treatment with DMSO, there is a shift in cells from SG2M to G1 (Figure 1C and 1D). We next used fluorescence activated cell sorting (FACs) to isolate cells from early G1, late G1, and SG2M phases from control and 24h DMSO-treated H9 FUCCI hPSCs (Figure 1B and 1D) and performed RNA-sequencing. Using principal component analysis (PCA), we found that the strongest source of variation was in treatment vs control (PC1), followed by phase of the cell cycle (PC2) (Figure 1E). We next used the nonparametric Dirichlet process Gaussian process mixture model (DPGP)^26^ to cluster fold changes in aligned reads (normalized to transcripts per million (TPM)) to assess changes in gene expression patterns associated with cell cycle progression. A total of 2972 differentially expressed genes (FDR < 0.05) underwent clustering and 10 clusters emerged (Figure 1F) with genes upregulated or downregulated in response to phase of the cell cycle following the DMSO treatment. The largest clusters consisted of genes with decreased expression in late G1 but high in early G1 and SG2M (cluster 7 with 454 genes) or increased expression in late G1 and reduced in early G1 and SG2M (cluster 5 with 420 genes) following the DMSO treatment. Genes with trajectories characteristic of the 10 clusters include those playing important roles in early development and regulating growth signaling pathways (e.g. PGAM1, LEFTY2, RHOB, WNT3, PIK3R3), ubiquitination and DNA repair (CUL4A), DNA replication licensing (e.g. MCM3), maintaining cell shape and cytoskeletal interactions (e.g. VIM, RHOB), and regulating transcription, splicing, and translation of genes through critical RNA helicases and polymerases (e.g. DDX46, POLR2H) (Figure 1G). Annotation of the genes sets representative of each cluster using the Molecular Signatures Database (MSigDB) shows the most significant pathways enriched in the 10 clusters (Figure 1H). Across all clusters, the DMSO treatment targeted pathways known to be tightly coordinated with the cell cycle, playing critical roles in cytoskeletal organization and membrane structure, transcriptional regulation, cell growth control, and development (e.g. Rho GTPases signaling, mitochondrial biogenesis, rRNA processing, neddylation, protein folding, extracellular matrix organization, cilium assembly, Pre-mRNA processing, spliceosome) (Figure 1H). Genes associated with 5 of the 10 clusters (R5, R6, R8, R9, and R10) were enriched in the Processing of Capped Intron-Containing Pre-mRNA (Figure 1H), indicating an important role for the DMSO treatment in regulating the efficiency and fidelity of gene expression^27^. Many pathways associated with mitochondrial function were also enriched (clusters R5 and R8), consistent with recent work demonstrating that mitochondrial dynamics play critical roles in the developmental potential of hPSCs^28^. Overall, this data illustrates that the DMSO treatment changes the expression of these genes in a phase-specific manner in hPSCs and thereby restricts their activity in a temporal manner that is otherwise not present in the cell cycle of untreated control hPSCs.

**Figure 1.**
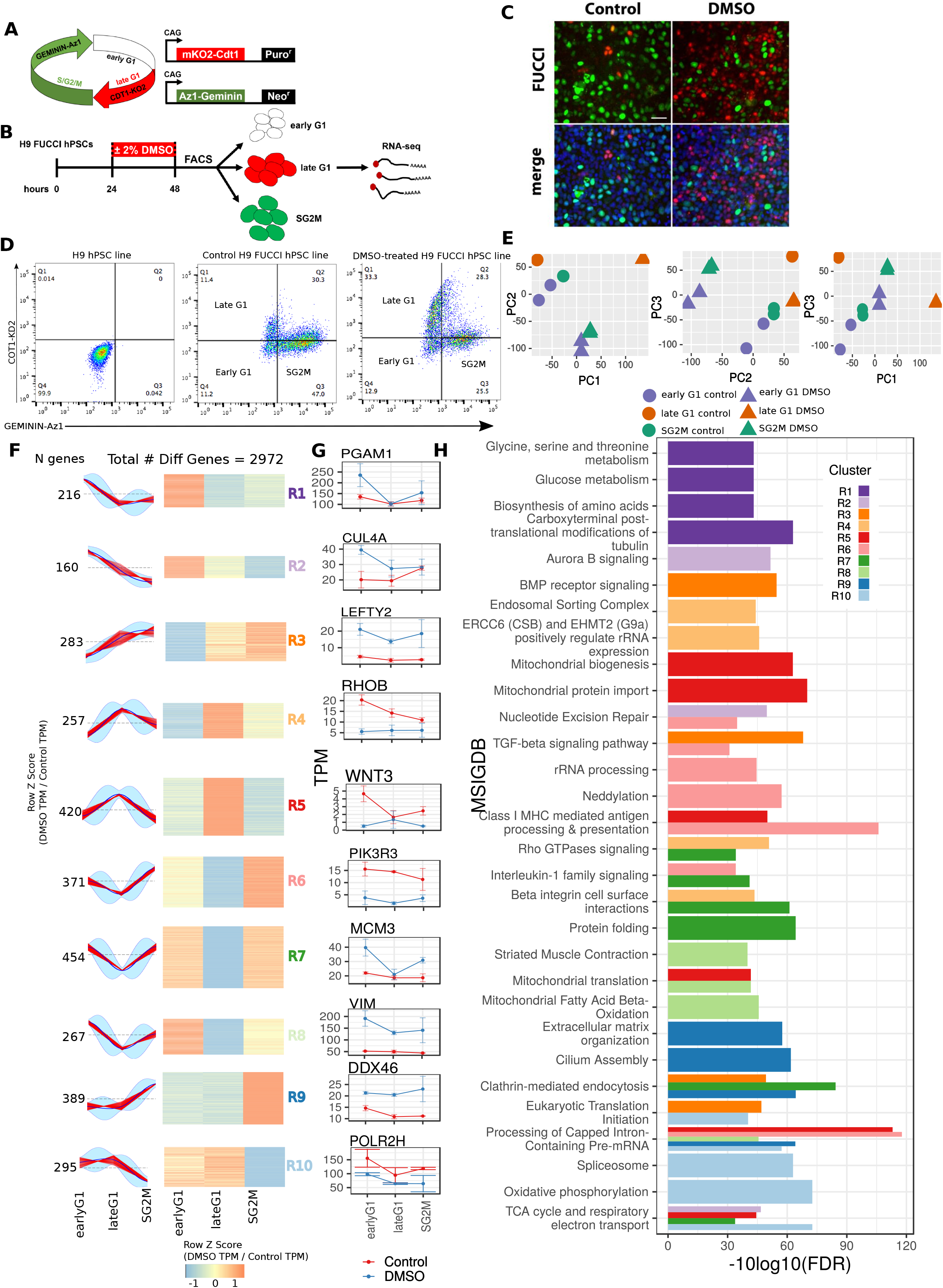
DMSO treatment of hPSCs changes gene expression trajectories in response to phase of the cell cycle. (A) Schematic representation of the FUCCI technology labeling individual G1 phase nuclei in red and S/G2/M phase nuclei in green. (B) Schematic of H9-FUCCI hPSCs treated with or without 2% DMSO for 24h followed by cell cycle sorting and high throughput RNA-sequencing. (C) Immunofluorescent images of Control and DMSO-treated H9 FUCCI hPSCs in the G1 phase (red) and SG2M phases (green) of the cell cycle. (D) Fluorescence activated cell sorting of cells in the early G1 (double negative), late G1 (red), and SG2M (green) phases of the cell cycle in control and DMSO-treated FUCCI hPSCs. (E) Principal component analysis of batch-corrected RNA-seq expression data. PC 1 (97.02% variance explained) vs PC 3 (0.38% variance explained) are plotted on TPM (transcript per million) data for hg19. (F) DPGP (Dirichlet process Gaussian mixture model) clustering of differentially expressed genes. DPGP clustering was applied to the fold change of (TPM DMSO / TPM control) for differential genes, and yielded 10 clusters, labeled R1 - R10. The Z-scores for genes in each cluster are plotted in heatmap form as well as line plots of trajectories across early G1, late G1, SG2M. Red lines indicate fold change trajectories for individual peaks assigned to the cluster. The light blue cloud indicates values within 2 standard deviations of the cluster mean. (G) Representative differentially expressed genes for each DPGP cluster R1-R10. TPM values with standard error bars are indicated for Control (red) and DMSO (blue) at the early G1, late G1, and SG2M phases. (H) Most significant MSIGDB pathways enriched in the DGPGP clusters R1 - R10. Height of the bar indicates -log10(FDR) values for the corresponding clusters.

### Regulatory role for PI3K-AKT signaling in hPSC differentiation

In aggregate, UpSet analysis shows that the most number of differentially expressed genes occur in the late G1 phase and are specific to distinct phases of the cell cycle -- of the 1078 genes downregulated in late G1, 783 were not significantly altered at the other cell cycle phases; of the 895 upregulated genes, 554 were unique to late G1 (Figure 2A). MSIGDB pathway analysis (FDR < 0.01) shows that DMSO affects a number of pathways associated with cell signaling (Figure 2B). Across all of the signaling pathways targeted by DMSO, PI3K was the most commonly represented gene followed by PIK3CA, a catalytic subunit of PI3K (Figure 2B). Kyoto Encyclopaedia of Genes and Genomes (KEGG) analysis shows that 48 genes associated with the PI3K-AKT pathway are significantly regulated by the DMSO treatment at one or more phases of the cell cycle (Figure 2C, Supplementary Figure 1). Many genes upstream in the pathway (e.g. PI3K receptors, PI3K, Ras) are generally down-regulated in the early and late G1 phases of the cell cycle. Other signaling pathways regulated by DMSO also converge upon PI3K and PI3KR signaling (examples illustrated in Supplementary Figures 2, 3, and 4), a pathway well known to regulate cell cycle, proliferation, differentiation, apoptosis, and growth and metabolism^29–35^. Pathways and genes associated with mitosis and the cell cycle (e.g. cell cycle checkpoints, p-value=3.98e-8) were also significantly regulated by the DMSO treatment through MSIGDB pathway and gene ontology (GO) enrichment analyses (Supplementary Figure 6). Expression patterns for genes commonly implicated in cell division or regulating early differentiation of hPSCs ^6,7^ are shown for DMSO-treated hPSCs compared to untreated control hPSCs as cells progress through the cell cycle (Supplementary Figure 6). Interestingly, pluripotency genes (GO Term GO:0019827 Pluripotency Genes; FDR=8.50e-1 by Fischer’s Exact Test) were not altered, suggesting that the DMSO effect on improved differentiation is not mediated by altering the expression of the pluripotency network. Given the convergence towards PI3K, we next investigated whether inhibiting PI3K would mimic the DMSO treatment and increase the mutlilineage differentiation potential of hPSCs. To suppress PI3K signaling, we treated H9 hPSCs with small molecule PI3 kinase inhibitors (LY294002 and Wortmannin) for 24h and subsequently induced differentiation into the ectodermal, mesodermal, and endodermal germ layers using previously published protocols (Figure 2D). Following directed differentiation, protein expression of germ layer specific genes^36^, Sox1 (ectoderm), Brachyury (mesoderm), and Sox17 (endoderm) were assessed by immunostaining. Treatment with the PI3K inhibitors increased subsequent differentiation capacity across all germ layers in a dose-dependent manner (Figure 2E and 2F). Similar improvements in differentiation were observed in another hPSC line, HUES6, known to have a very poor propensity for differentiation^4^ (Supplementary Figure 5). Together, these results show that understanding gene trajectories in the cell cycle of hPSCs can highlight important signaling mechanisms regulating hPSC differentiation.

**Figure 2.**
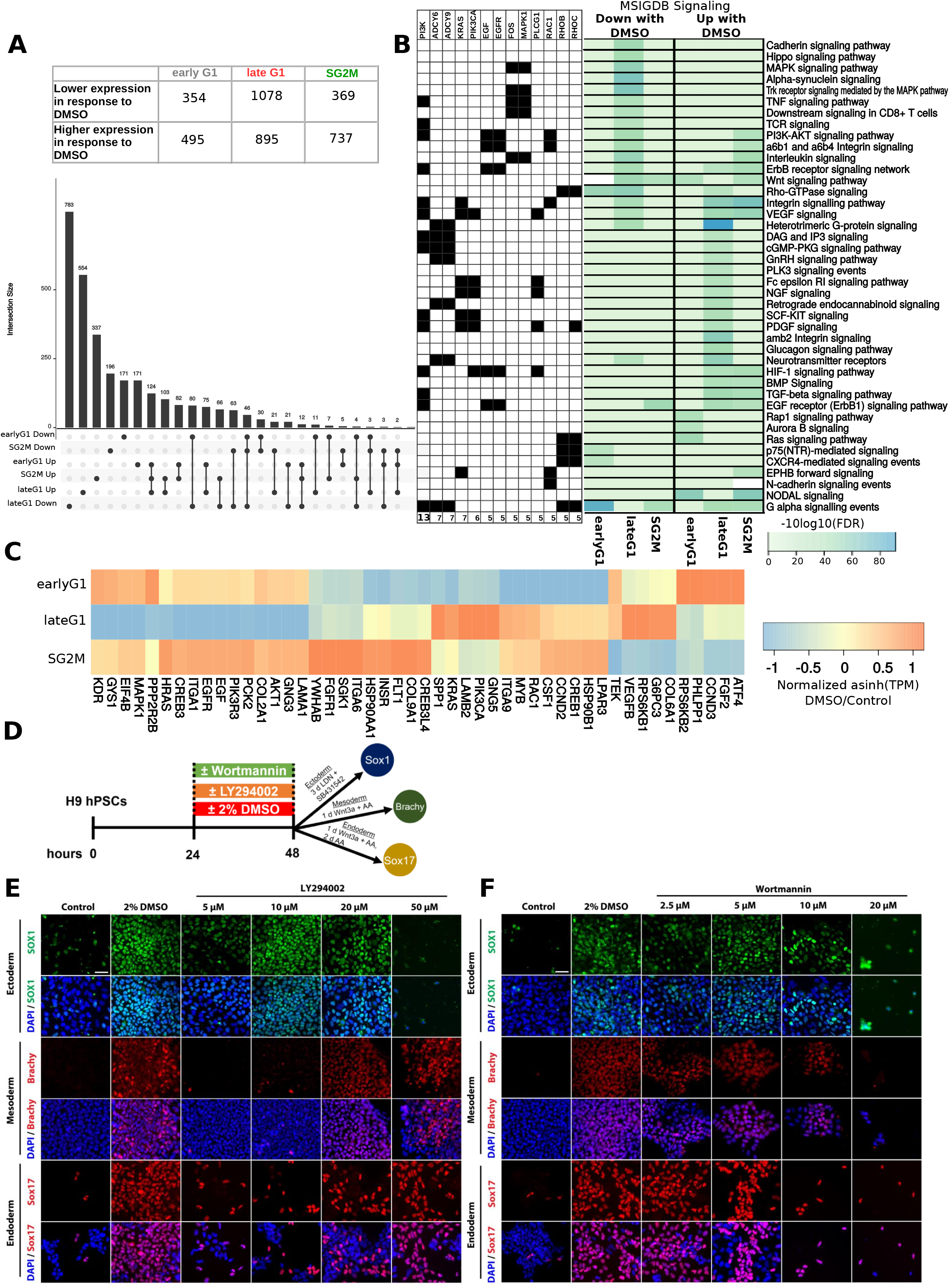
PI3K inhibition increases hPSC differentiation across all germ layers. (A) Number of differentially expressed genes (FDR < 0.05, LFC>= 1) in Control vs DMSO-treated FUCCI hPSCs in the early G1, late G1, and SG2M phases. UpsetR diagram of differentially expressed genes shows that number of differential genes that increase (up) or decrease (down) in expression in response to DMSO at the early G1, late G1, and/or SG2M phases. (B) Enriched REACTOME pathways for differential genes at the early G1, late G1, and SG2M phases of the cell cycle. The heatmap shading corresponds to the -10log10(FDR) for each pathway across the different phases of the cell cycle. Genes with differential expression in response to DMSO treatment that are present in five or more differential signaling pathways are indicated with black boxes in the grid to the left of the heatmap. (C) Differentially expressed genes within the PI3K-AKT signaling pathway. Heatmap values are row z-scores of asinh(TPM) DMSO / asinh(TPM) controls. (D) Schematic of H9 hPSCs treated with 2% DMSO or inhibitors of PI3K (LY294002 or Wortmannin) for 24 hours and subsequently directly differentiated into the ectodermal, mesodermal, and endodermal germ layers. Immunostaining for germ layer specific markers following treatment with (E) LY294002 or (F) Wortmannin compared with untreated Control and 2% DMSO-treated hPSCs.

## DISCUSSION

Strikingly, although DMSO is an agent with pleiotropic effects^37,38^, here, we show that a short 24h treatment of hPSCs targets 2,972 genes in an orchestrated manner, particularly those controlling cell division and early developmental pathways. Genes are periodically expressed because there is special need for the gene products at particular points in the cell cycle^39^. Genes associated with cytoskeletal, cilium assembly, and cell adhesion factors were especially subject to regulation by the DMSO treatment in the SG2M phases, characteristic of a time when cells may need to duplicate centrioles in the S phase, change shape during mitosis, or exit the mitotic cycle to differentiate^40^. Many of the targeted pathways play critical roles during embryogenesis, including Wnt, BMP, NODAL, FGF, Hippo, EGF, VEGF, and PDGF as well as the downstream signaling pathways such as MAPK, Trk receptor, and PI3K^41^. Integration of these signaling pathways coordinates a number of developmental processes, including proliferation, fate determination, differentiation, apoptosis, migration, adhesion, and cell shape, to ultimately affect organogenesis. Most of the pathways that were regulated by DMSO converged on PI3 kinase signaling. Concordantly, suppression of PI3K signaling increased differentiation propensity across all germ layers in hPSCs, highlighting the utility of the genome-wide profiling approach used here to dissect out important signaling mechanisms regulating the developmental potential of pluripotent stem cells. This work is consistent with prior studies showing that PI3K-dependent signals promote embryonic stem cell proliferation and supports the notion that each phase of the cell cycle is important in performing distinct roles to orchestrate stem cell fate^42,43^. Many of the signaling pathways and effects on metabolic function and cell adhesion identified here were also reported to play important regulatory roles during early transitions in pig embryonic development in recent work^44^, suggesting shared mechanisms across species.

It would be interesting to investigate if improvements in terminal differentiation or enhancements in CRISPR-mediated genome editing of hPSCs following a 24h DMSO treatment^45^ may be due to changes in the molecular properties elicited on the pluripotent cell cycle. In conclusion, our data yield novel insights on the transcriptional and signaling dynamics during early transitory states in human pluripotent stem cells that could be a useful point of focus in studying embryonic development. Targeting these early modes of regulation may put hPSCs on a better trajectory for differentiation and ultimately improve their utility for regenerative medicine.

## Acknowledgments

We are very grateful to Stephen Dalton and Amar Singh for providing the H9 FUCCI cell line, Beijing Wu and Trent Edwards for their excellent technical support with experiments, and the Flow Cytometry Core facilities for their assistance. We thank Thomas Brickler, Jing Bian, Danielle Sambo, and Liang Ma for their feedback on the manuscript. This work was supported by grants from the Stanford University School of Medicine, a Siebel Fellowship awarded to S.C., and the US National Institutes of Health (S10OD018220).

## Materials and Methods

### hPSC maintenance conditions

All hPSC cultures were maintained at 37°C, 5% CO2 and expanded on Matrigel (BD Biosciences) coated plates in mTeSR (STEMCELL Technologies) with 10 μM ROCK inhibitor Y-27632 (Abcam). The previously characterized human embryonic stem cell lines (HUES6, H9 (Wicell), H9 FUCCI) were used in this study^4,46^. The H9 FUCCI hPSC line was constructed by Singh et al., 2013 by introducing fluorescent reporters into expression vectors under the control of the constitutive CAG promoter and linked to neomycin or puromycin selectable markers through an internal ribosome entry site^7,47^. G418 sulfate and the puromycin were used to maintain the H9-FUCCI hPSCs.

### Regulatory and institutional review

All human pluripotent stem cell experiments were conducted in accord with experimental protocols approved by the Stanford Stem Cell Research Oversight (SCRO) committee.

### DMSO treatment and differentiation protocols

For all RNA-seq experiments, H9 FUCCI hPSCs were plated onto plates coated with growth factor–reduced Matrigel (BD Biosciences) in mTeSR with 10 μM ROCK inhibitor Y-27632 (Abcam) at 1 million cells per well of a 6-well plate. After 24h, cells were cultured in mTeSR with or with 2% DMSO for another 24h. After a 24h DMSO treatment, cells were collected and prepared for fluorescence-activated cell sorting by flow cytometry for subsequent RNA-seq analyses.

For differentiation experiments, after a 24h treatment with DMSO or the PI3K inhibitors LY294002 (Selleck Chemicals) and Wortmannin (Selleck Chemicals) in mTESR, the medium was replaced with one of the following media at the start of each differentiation, with media replacement every day in all protocols.

#### Ectoderm

Ectoderm differentiation was induced using Knockout DMEM (Life Technologies), LDN-193189 (100 nM; Stemgent) and SB431542 (10 µM; Tocris) containing 15% Knockout serum replacement (Life Technologies) for 3 days. The medium was removed and replaced with fresh medium every 24 hours.

#### Mesoderm

Mesoderm differentiation was induced in advanced RPMI medium (Invitrogen) supplemented with Wnt3a (20 ng/ml; R&D Systems) and Activin A (100 ng/ml; R&D Systems) for 24h.

#### Endoderm

Endoderm differentiation was induced in RPMI medium (Invitrogen), supplemented with Wnt3a (20 ng/ml; R&D Systems) and Activin A (100 ng/ml; R&D Systems) for 24 hours and subsequently in RPMI medium containing Activin A (100 ng/ml) for 2 days.

### Immunocytochemistry

Cells were rinsed in PBS and fixed in 4% paraformaldehyde (PFA; Sigma) for 30 min. Following the rinses, cells were blocked for 1 h at room temperature in 5% donkey serum (Jackson ImmunoResearch), 0.3% Triton X-100 in PBS. All primary antibody incubations were done overnight at 4°C in blocking solution at a 1:500 dilution unless otherwise noted. Primary antibodies used in this study were: Sox1 (R&D Systems), Brachyury (R&D Systems), and Sox17 (R&D Systems). Cells were rinsed the next day, followed by secondary antibody incubation for 1 h at room temperature at a 1:500 dilution. Secondary antibodies (Invitrogen) conjugated to Alexa Fluor 488 or 594 were used to visualize primary antibodies. DAPI (4,6-diamidino-2-phenylindol, Life Technologies) was used as a nuclear dye to stain all cells. All images were acquired using a Leica Fluorescent Microscope.

### Fluorescence-activated cell sorting (FACS)

Following a 24h treatment with or without 2% DMSO, the H9-FUCCI hPSCs were collected and resuspended in FACS buffer (0.2% BSA in PBS). Cells in late G1 phase express mKO2-Cdt1 (color red), while the S/G2/M cells express mAG-Gem (color green); the double-negative population is indicative of early G1 cells. DAPI was included to stain and gate out the dead cell population. The H9 hPSC line was also included as a control and analyzed by flow cytometry. Cells were run on a flow cytometer adjusted for UV excitation to measure DAPI fluorescence at blue wavelengths and RFP and GFP wavelengths to isolate and sort populations from the early G1, late G1 and SG2M fractions of the cell cycle. 50,000 cells were collected for each population in replicate over two independent experiments.

### RNA-sequencing library preparation

For bulk-population RNA-seq, RNA was extracted from cell cycle sorted control and DMSO-treated hPSCs. Integrity of extracted RNA was assayed by on-chip electrophoresis (Agilent Bioanalyzer) and only samples with a high RNA integrity (RIN) value were used for RNA-seq. Purified total RNA was reverse-transcribed into cDNA using the Ovation RNA-seq System V2 (NuGEN) and cDNA was sheared using the Covaris S2 system (duty cycle 10%, intensity 5, cycle/burst 100, total time 5 min). Sheared cDNA was cleaned up using Agencourt AMPure XP beads (Beckman Coulter) and ligated to adaptors (Illumina) Sequencing libraries were constructed using the NEBNext Ultra DNA Library Prep Kit (New England Biolabs) using barcoded adaptors to enable multiplexing of libraries on the same sequencing lane. For each RNA-seq library, the effectiveness of adaptor ligation and effective library concentration was determined by Bioanalyzer before loading them in multiplexed fashion onto an Illumina HiSeq4000 (Stanford Functional Genomics Facility) to obtain ∼150 bp paired-end reads.

### RNA-sequencing processing pipelines

Paired end Illumina RNA-seq reads were trimmed to 75 base pairs using the Trimommatic software^48^(v. 0.36) The trimmed reads were aligned to the hg19 reference genome using STAR (v2.5.3a)^49^. All parameters were set to their default values, with the exception of the following:

--outFilterScoreMinOverLread 0

--outFilterMatchNminOverLread 0

--outFilterMatchNmin 0

--outFilterMismatchNmax 2

These parameters were set less stringently relative to their default values to enable less stringent determination of multimappers, ultimately leading to an improved overall alignment rate. Gene and transcript expression was then quantified with RSEM(v1.1.17)^50^, also using the default parameters. Gene expression values were normalized as asinh(transcripts per million).

Differential gene expression was calculated on normalized read counts. TPM (transcript per million) values from RSEM were used for differential expression analysis. The sva package^51^ in R was used to perform surrogate variable analysis with the null model: expression ∼ cell cycle phase + DMSO treatment status. The significant non-correlated surrogate variables were identified and subtracted from the normalized data. The limma R package^52^ was used to identify differentially expressed genes with the model as mentioned above. Three sets of pairwise comparisons were performed with limma: early G1, DMSO vs early G1, Control; late G1, DMSO vs late G1, Control; SG2M, DMSO vs SG2M, Control. Differential genes were those with FDR values < 0.01 and abs(log2(fold change)) >1.

Differential genes underwent clustering with the Dirichlet process Gaussian process mixture model DPGP^26^ software. 1000 iterations of clustering were performed with default software parameters. Inputs to the clustering algorithm for genes consisted of the fold change in gene expression in DMSO samples over control samples for all differential genes, normalized as asinh(TPM - surrogate variable contribution). Differential gene sets were visualized with the UpSetR package^53^.

Pathway analysis (MSIGDB, KEGG) was performed with DAVID bioinformatics software (v. 6.8)^54^. Genes within each DPGP cluster were provided to DAVID as a gene list, with the hg19 reference gene list as background. Pathways with FDR values <0.01 were determined to be significant. The differential genes present in the top 50 most significant pathways (42 associated with signaling; 8 associated with mitosis) identified by the Cytoscape Reactome FI plugin were aggregated across pathways **(Figure 2B, Supplementary Figure 6B)**. The number of pathways that each gene was associated with was tallied. It was determined that the PI3K gene was present in 13 of the top 50 differential Reactome pathways, and this observation was confirmed by querying the PI3K gene in the WikiPathways database^55^ and cross-checking all pathways that were associated with the query.

## Figure Legends

**Supplementary Figure 1. Differentially expressed genes within the PI3K-AKT signaling pathway**

KEGG annotation for the PI3K-AKT signaling pathway. Genes with differential expression in DMSO-treated hPSCs vs control hPSCs are filled in gray. Genes downregulated or upregulated in response to DMSO treatment are denoted by the color-coded triangles when differential at the early G1, late G1, and/or SG2M phases.

**Supplementary Figure 2. Differentially expressed genes within the TNF signaling pathway**

(A) KEGG annotation for the TNF signaling pathway. Genes with differential expression in DMSO-treated hPSCs vs control hPSCs are filled in gray. Genes downregulated or upregulated in response to DMSO treatment are denoted by the color-coded triangles when differential at the early G1, late G1, and/or SG2M phases. (B) Summary of differentially expressed genes within the TNF signaling pathway. Heatmap values are row z-scores of asinh(TPM) DMSO / asinh(TPM) controls.

**Supplementary Figure 3. Differentially expressed genes within the cGMP-PKG signaling pathway**

(A) KEGG annotation for the cGMP-PKG signaling pathway. Genes with differential expression in DMSO-treated hPSCs vs control hPSCs are filled in gray. Genes downregulated or upregulated in response to DMSO treatment are denoted by the color-coded triangles when differential at the early G1, late G1, and/or SG2M phases. (B) Summary of differentially expressed genes within the WNT signaling pathway. Heatmap values are row z-scores of asinh(TPM) DMSO / asinh(TPM) controls.

**Supplementary Figure 4. Differentially expressed genes within the VEGF signaling pathway**

(A) KEGG annotation for the VEGF signaling pathway. Genes with differential expression in DMSO-treated hPSCs vs control hPSCs are filled in gray. Genes downregulated or upregulated in response to DMSO treatment are denoted by the color-coded triangles when differential at the early G1, late G1, and/or SG2M phases. (B) Summary of differentially expressed genes within the VEGF signaling pathway. Heatmap values are row z-scores of asinh(TPM) DMSO / asinh(TPM) controls.

**Supplementary Figure 5. PI3K inhibition increases HUES6 hPSC differentiation across all germ layers**

(A) Schematic of HUES6 hPSCs treated with 2% DMSO or inhibitors of PI3K (LY294002 or Wortmannin) for 24 hours and subsequently directly differentiated into the ectodermal, mesodermal, and endodermal germ layers. Immunostaining for germ layer specific markers following treatment with (B) LY294002 or (C) Wortmannin compared with untreated Control and 2% DMSO-treated hPSCs.

**Supplementary Figure 6. DMSO treatment regulates the cell cycle of hPSCs**

(A) TPM values for cell-cycle associated genes are illustrated for DMSO-treated hPSCs (blue) and control hPSCs (red) at the early G1, late G1, and SG2M phases of the cell cycle. * denotes FDR < 0.05. (B) Enriched REACTOME pathways for differential genes associated with Mitosis at the early G1, late G1, and SG2M phases of the cell cycle. The heatmap shading corresponds to the -10log10(FDR) for each pathway across the different phases of the cell cycle. (C) Fold change row z-scores of asinh(tpm) DMSO/control for differentially expressed genes that are associated with enriched sub-terms of the cell cycle biological process GO Term (GO:0007049). (D) -10log10(FDR) for enriched GO terms associated with the cell cycle.

## Table Legends

**Supplementary Table 1**

Data QC for paired-end Illumina RNA-seq samples, including input number of reads as well as percent of uniquely mapped reads.

