## Supplementary figures and images for "Cell cycle dynamics of human pluripotent stem cells primed for differentiation"

### Supplementary Figure 1

#### Supplementary Figure 1

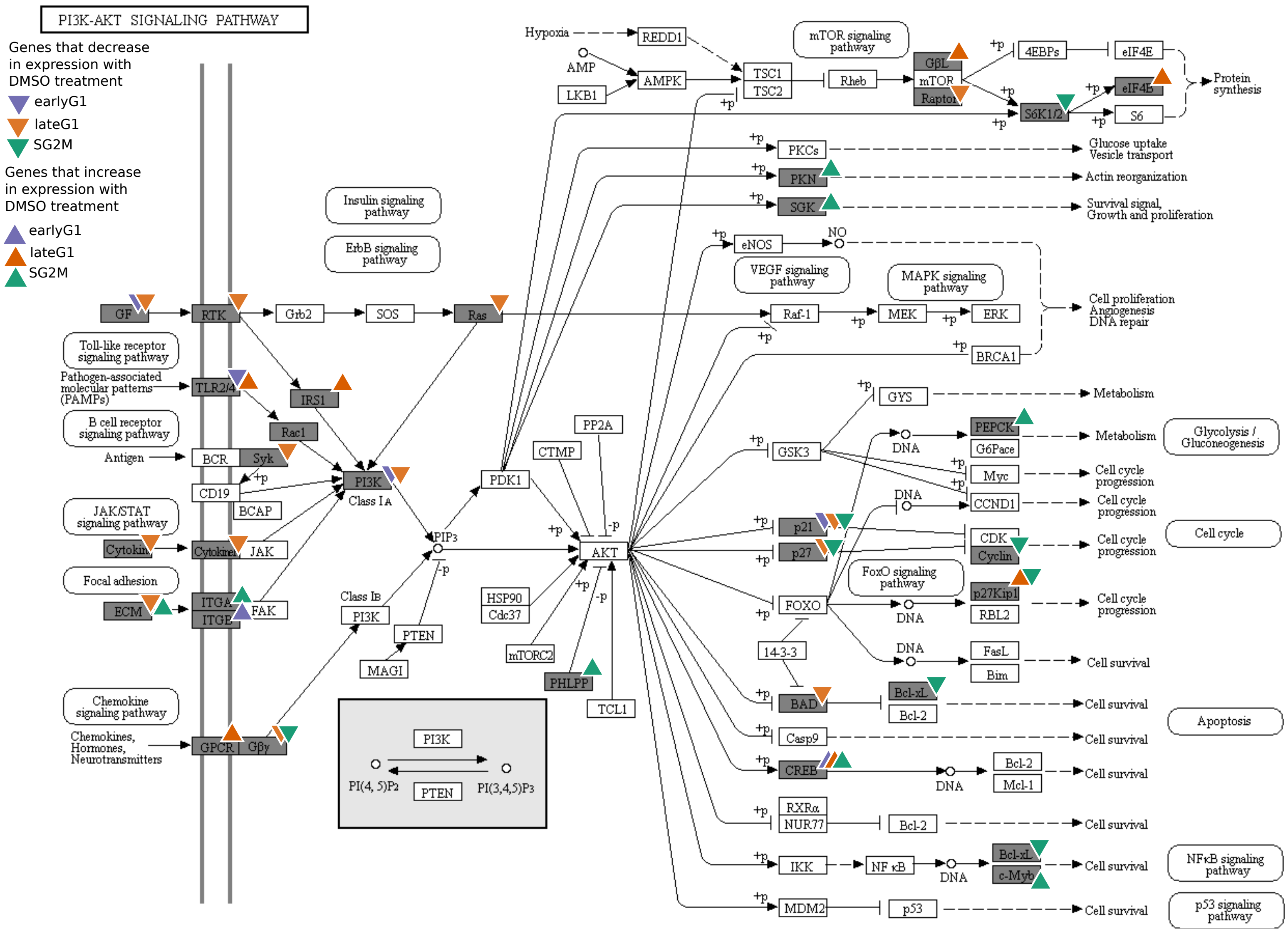

### Supplementary Figure 2

#### A Supplementary Figure 2

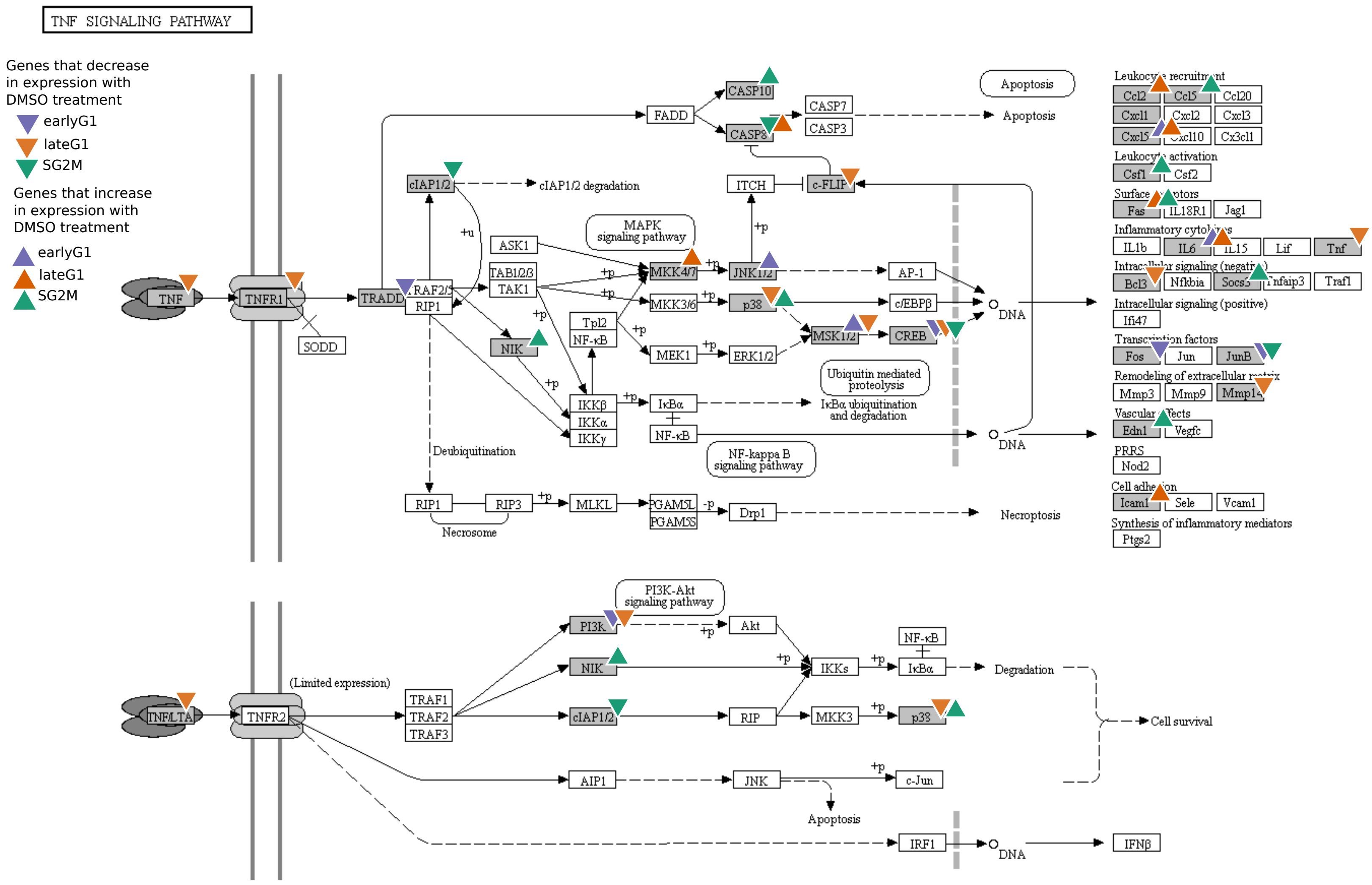


**B**

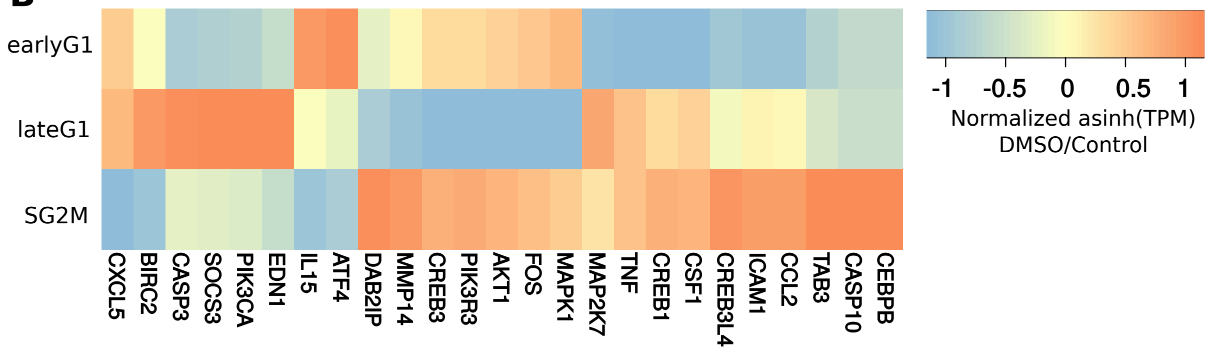

### Supplementary Figure 4

#### A Supplementary Figure 4

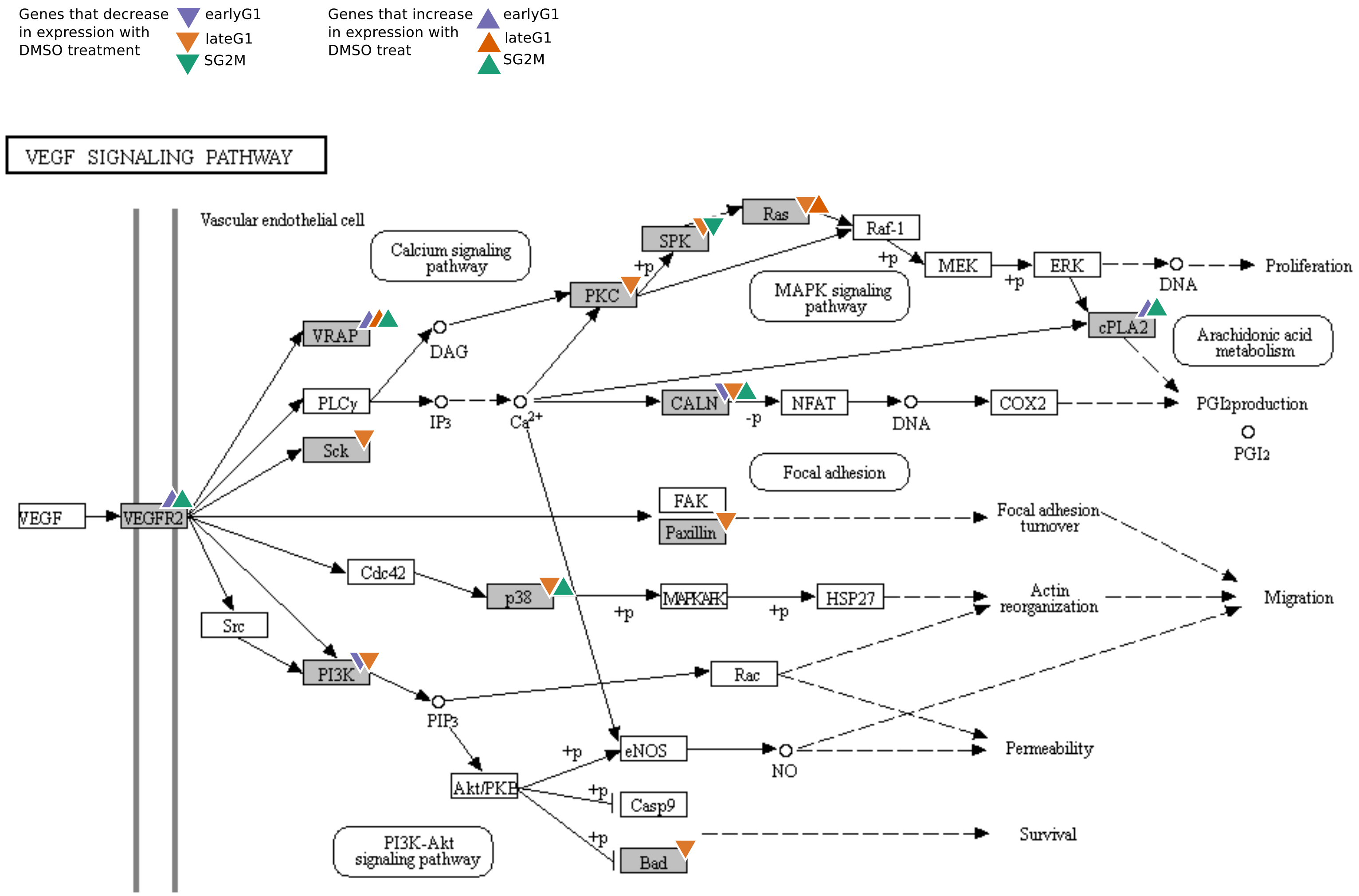


**B**

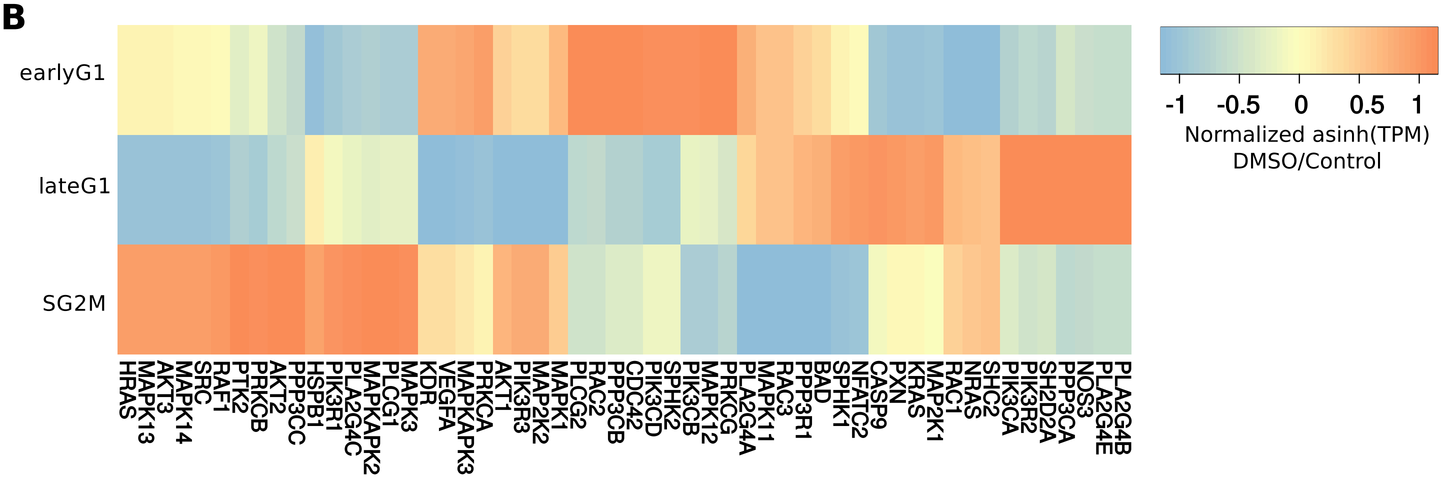

### Supplementary Figure 5

Supplementary Figure 5

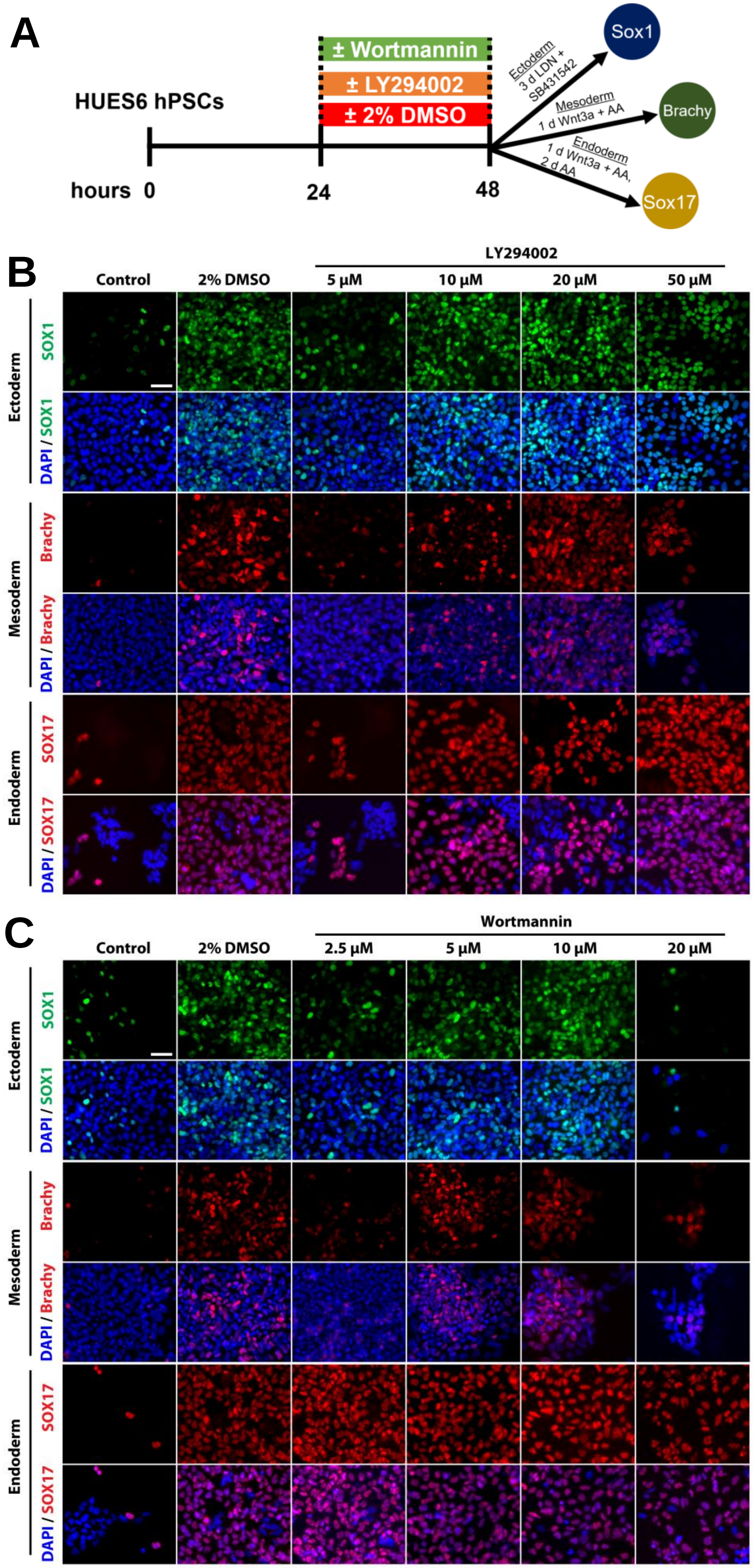

### Supplementary Figure 6

#### Supplementary Figure 6

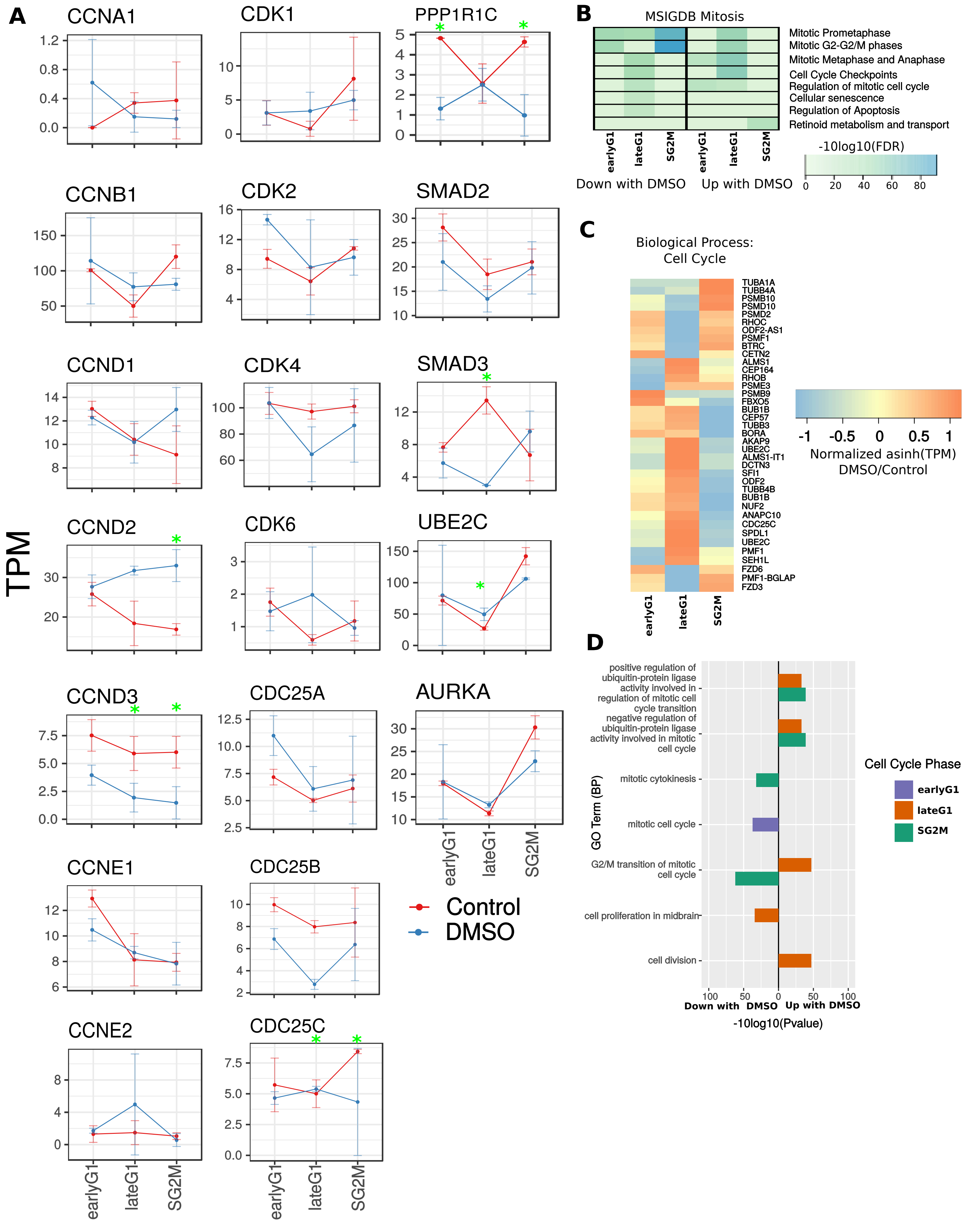
