## Supplementary Figure 3 for "Cell cycle dynamics of human pluripotent stem cells primed for differentiation"

A

Genes that decrease in expression with DMSO treatment: earlyG1, lateG1, SG2M

Genes that increase in expression with DMSO treat: earlyG1, lateG1, SG2M

B

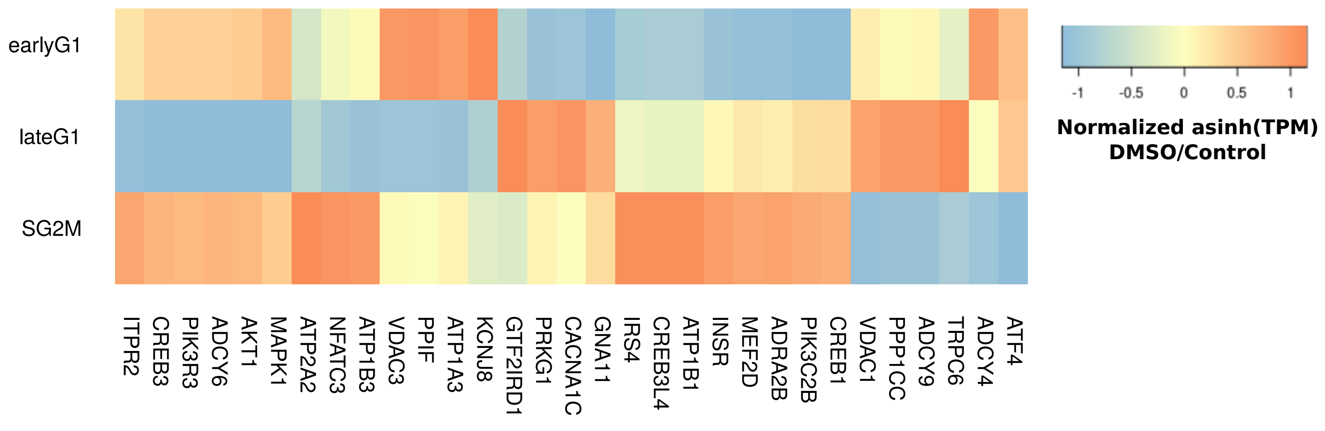
